# NL2BLTL: Automated Generation of Bounded Linear Temporal Logic from Natural Language for Systems Biology

**DOI:** 10.1101/2025.08.06.668950

**Authors:** Difei Tang, Natasa Miskov-Zivanov

## Abstract

**Motivation:** Translating natural language biological discoveries into formal temporal logic specifications for model verification demands expertise most experimentalists lack. Bounded Linear Temporal Logic (BLTL), which attaches explicit time bounds to temporal operators, is well suited for capturing biological dynamics. However, no automated solution exists for this translation, limiting the broader adoption of statistical model checking in systems biology.

**Results:** We present NL2BLTL, the first framework to automate natural language to BLTL translation for systems biology. NL2BLTL combines a synthetic dataset of 5,000 NL-BLTL pairs built via grammar- guided generation, Chain-of-Thought preprocessing to resolve linguistic ambiguity in biological hypotheses, and grammar-constrained decoding to enforce syntactic validity and prevent hallucinated variables or time bounds. Evaluated on a newly curated biomedical dataset drawn from published T cell and pancreatic cancer models, NL2BLTL framework achieves 84.62% exact match and 100% syntactic validity, outperforming GPT-4 by over 16 points and improving 14 points over the base fine-tuned model.

**Availability and Implementation:** NL2BLTL and both datasets are freely available at https://github.com/pitt-miskov-zivanov-lab/BioBLTL.

## 1 Introduction

An important challenge in computational systems biology is the verification of computational model behavior against experimental observations or hypotheses. Expressing these biological propositions formally enables a rigorous assessment of model behavior through automated verification techniques such as model checking^1^.

Bounded Linear Temporal Logic (BLTL)^2^ is among the most expressive and practically useful formalisms for verifying whether a computational model reproduces experimentally observed dynamics. Unlike general temporal logics, BLTL allows modelers to attach explicit numerical time bounds to temporal operators, making it uniquely suited to the time-sensitive dynamics of biological systems, from the activation and inhibition of signaling cascades to the stabilization of cell fate decisions. Statistical model checking^3^ with BLTL can systematically evaluate whether a model’s simulated trajectories satisfy a given specification, providing quantitative confidence estimates rather than simple qualitative pass/fail results. However, efficient translation of biological hypotheses expressed in natural language (NL) into valid BLTL formulae requires knowledge of formal syntax and semantics that lies outside the training of most experimental biologists. This gap could be overcome through efficient automated translation, yet such methods are still lacking.

Early approaches to natural language to temporal logic (NL-TL) translation relied on template matching^4,5^, semantic parsing^6^, rule-based grammars^7^, or controlled natural languages to generate formalisms such as Linear Temporal Logic (LTL)^8^, Computation Tree Logic (CTL)^9^, and Signal Temporal Logic (STL)^10^. More recent work^11,12^ explored multi-stage pipelines that first convert natural language into an intermediate representation using large language models (LLMs) and then map that representation to temporal logic formulas. While effective in constrained settings, these approaches have limited generalizability and flexibility in real-world applications. For example, NL2TL^13^ leverages LLMs to construct a synthetic NL- TL dataset, which is then applied to fine-tune T5^14^, a pre-trained sequence-to-sequence language model. On the other hand, KGST^15^ first uses LLMs fine-tuned on an NL-STL dataset to generate preliminary STLs, and then evaluates and refines them using GPT by comparing each sample with retrieved NL-STL pairs as external knowledge. More recently, another LLM-based approach^16^ translates biomedical observations into STL specifications using structured syntactic and semantic feedback.

Furthermore, while several studies have explored NL-TL translation for software verification^17^ and cyber-physical systems^18,19^, biological systems present distinct challenges these approaches do not address: temporal cues in biological hypotheses are often implicit, numerical time bounds are embedded in experimental context, and species names and activation states must match precisely those defined in the model. No framework has been designed specifically for this problem.

We present NL2BLTL, the first dedicated framework for automated natural language to BLTL translation in systems biology, designed to overcome three core obstacles. Annotated NL-BLTL training data is notably lacking in biology, so we construct a synthetic dataset of 5,000 NL-BLTL pairs through a grammar- guided generator grounded in biological species from the Cell Collective database^20^, paired with GPT-4^21^- driven natural language synthesis. Further, biological text is inherently ambiguous in temporal expression, so we introduce a Chain-of-Thought (CoT)^22^ preprocessing step that decomposes each input into explicit temporal and logical components before translation. Finally, LLMs are prone to hallucination, generating syntactically invalid formulas or introducing variables and time bounds absent from the input, so we apply grammar-constrained decoding (GCD)^23^ that restricts generation to the defined BLTL grammar using constraints extracted during CoT preprocessing.

The contributions of this work are fourfold. First, NL2BLTL is, to our knowledge, the first framework for NL-to-BLTL transformation in systems biology, making formal verification accessible to experimental biologists without requiring them to learn temporal logic. Second, we define a set of structural templates that capture the recurring temporal patterns observed in curated BLTL specifications from prior systems biology modeling studies. These templates form the basis of our synthetic data generator and serve as reusable specification patterns for BLTL-based modeling and verification in this work. Third, we release two new NL-BLTL datasets, a large-scale synthetic NL-BLTL dataset for training and a manually curated biomedical dataset drawn from published T cell differentiation and pancreatic cancer cell models, establishing the first evaluation resource for this task. Fourth, fine-tuned open-source LLMs within our framework substantially outperform few-shot GPT-4, with Qwen2.5 achieving 84.62% exact match and 100% syntactic validity on the biomedical dataset. Ablation studies confirm that CoT and GCD make complementary contributions, together accounting for a 14-point gain in exact match over the base fine-tuned model, providing clear design guidance for future work in translating natural language into formal biological specification.

## 2 Methods

Figure 1 presents an overview of our NL-BLTL generation framework. Our approach consists of three main components: synthetic data generation, model fine-tuning, and inference using CoT-based NL preprocessing and GCD. Prior to data generation, we performed a structural pattern analysis of curated BLTL specifications in systems biology to identify recurring temporal structures.

**Figure 1.**
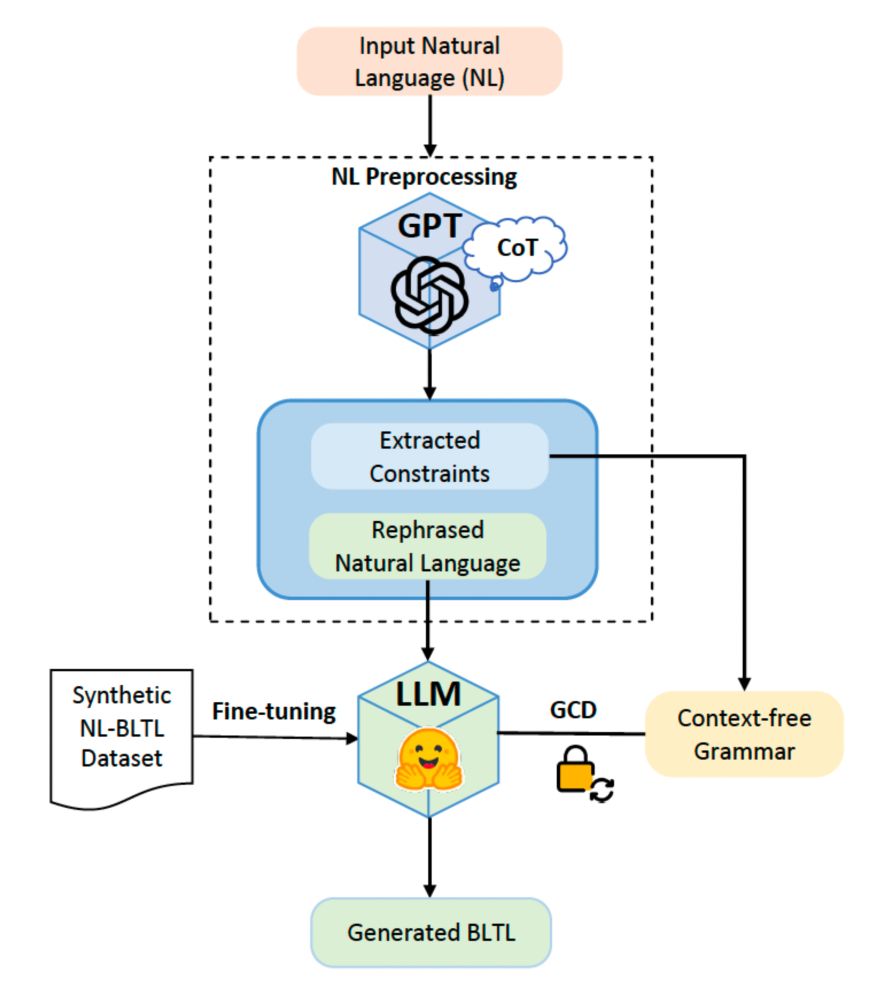
An overview of our framework for NL-BLTL transformation. An input NL description is preprocessed with CoT prompting to extract temporal and logical constraints. A fine-tuned LLM then generates the target BLTL under grammar-constrained decoding (GCD) using a context-free grammar augmented with the extracted constraints.

### 2.1 BLTL preliminaries

The base unit of a temporal logic formula is an *atomic proposition*, which takes the form x∼c, where x ∈ X and X is a set of variables representing system components or species, c is a numerical constant, and ∼ ∈ {>, ≥, <, ≤, =} is a relational operator.

In BLTL, atomic propositions are combined using two groups of operators. The first group includes Boolean operators ¬ (negation/not), ∧ (conjunction/and), and ∨ (disjunction/or), which combine atomic propositions into Boolean expressions. The second group consists of bounded temporal operators F[t] (future), G[t] (globally), and U[t] (until), each parameterized by a time bound t ∈ ℕ^+^ to allow specifications to refer to bounded futures in time.

A *formula* in BLTL is defined as:

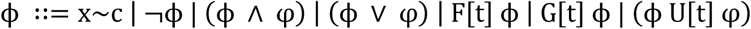

where **φ** and φ are BLTL formulas, and the base case x∼c is an atomic proposition as introduced above. The grammar is recursive such that each occurrence of **φ** or φ on the right-hand side denotes any BLTL formula derivable from this rule.

### 2.2 Systems biology preliminaries

In our setting, a biological model is associated with a finite set X of variables representing states of biological species. The explicit time bounds on temporal operators distinguish BLTL from LTL and make it particularly suitable for modeling time-dependent behaviors in biological processes. To match the concrete syntax accepted by model-checking tools, throughout the remainder of this paper we will write ‘&’ for ∧ and ‘|’ for ∨ in all BLTL formulas, grammar rules, and figures.

BLTL specifications are evaluated against simulation traces^24^ of elements in a computational model. A *trace* represents a sequence of states through which a species transitions over time. Given a BLTL specification and a set of traces, statistical model checking^3^ returns a probability estimate that quantifies how well the model’s behavior aligns with the specified dynamics.

To obtain biologically grounded species names for instantiating atomic propositions during synthetic data generation, we draw from the Cell Collective database^20^, which provides a large repository of curated models of cellular signaling and regulatory networks with consistent species naming. Cell Collective serves only as a source of species names in our pipeline and does not itself contain BLTL specifications. The curated BLTL specifications used elsewhere in this work are obtained from separate sources described in Sections 2.4 and 3.1.

### 2.3 Restricted BLTL-based grammar

We define here BLTL_R_, a restricted fragment of BLTL that captures expressive temporal patterns commonly observed in systems biology, including state transitions and stabilization, while ensuring that every generated formula is syntactically valid and compatible with statistical model-checking approaches^2,25^. As described in the following, BLTL_R_ is a subset of the set of all BLTL formulas derivable from the context-free grammar shown in Figure 2(a).

**Figure 2.**
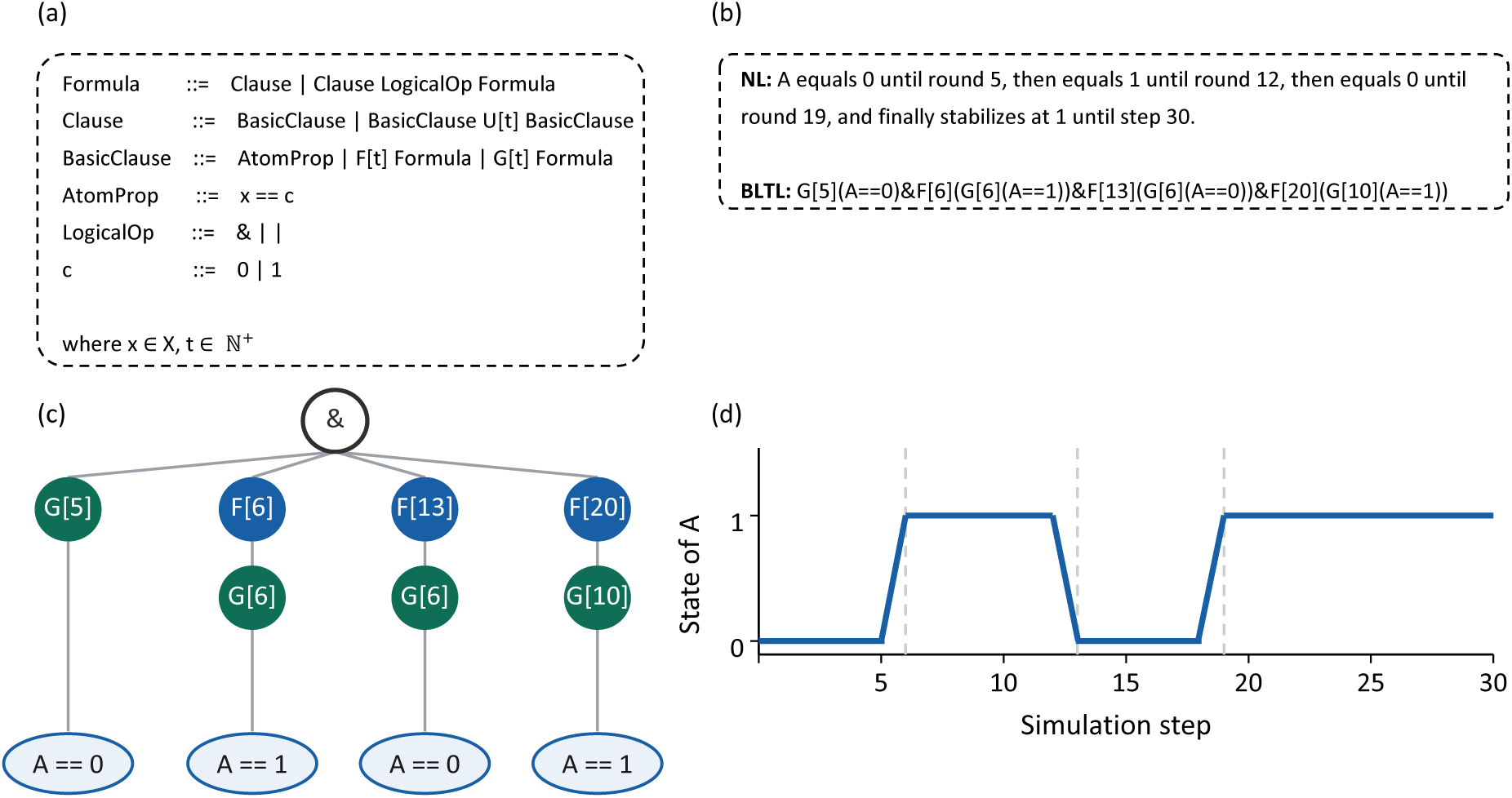
An example of a BLTL formula and its representations. (a). The context-free grammar defining BLTL_R_. AtomProp denotes an atomic proposition of the equality form x == c, where x is a species variable and c a numerical constant; F[t], G[t], U[t] are bounded temporal operators with time bound t. & and | are Boolean conjunction and disjunction. (b). A BLTL_R_ specification describing a temporal sequence of state transitions for species A, paired with its natural-language description. (c). The parse tree of the formula in (b). Internal nodes are temporal operators and Boolean connectives, and leaves are atomic propositions. (d). A simulation trace of A satisfying the formula, with shaded regions indicating the time intervals covered by each top-level subformula.

The grammar is built on three non-terminals: BasicClause, Clause and Formula. A BasicClause is either an atomic proposition (AtomProp) defined in Section 2.1 or a bounded temporal expression by applying F[t] or G[t] to Formula. A Clause consists of a BasicClause optionally extended with a bounded Until operator. A Formula consists of a finite number of Clause expressions connected by logical operators ‘&’ (and) or ‘|’ (or). We note here that negation is not included, and therefore, Formula is a restricted version of the general BLTL formula (Section 2.1). The grammar is defined recursively, allowing temporal operators to be applied to Formula expressions, which then enables the construction of nested temporal patterns such as sequential state transitions and stabilization. Atomic propositions in BLTL_R_ take the equality form x==c. Additionally, by construction, the algorithm limits the depth of nesting of temporal operators, and it also does not allow for the G[t] (F[d]**φ**) order (Section 2.4).

Figure 2(b-d) illustrates a BLTL_R_ specification of an example temporal behavior of species A, with its parse tree and the corresponding simulation trace. The natural language counterpart uses implicit temporal cues such as “then” and “stabilizes” that must be resolved into time-bound intervals, which is performed automatically by our CoT-based preprocessing step described in Section 2.7.

### 2.4 Structural templates of BLTL_R_ specifications

To characterize the temporal behaviors typically modeled in biological systems and to inform the design of our synthetic data generator, we analyzed 117 manually curated BLTL specifications and their corresponding NL descriptions, drawn from prior studies on naïve T cells^26^ and pancreatic cancer cell (PCC) microenvironment^25^. Despite substantial variability in biological contexts and linguistic expressions, the corresponding BLTL formulas include recurring structural patterns. We group these patterns into six representative template categories, which define the conceptual basis for the BLTL_R_ rules used in synthetic data generation (Section 2.5). Figure 3 summarizes these categories and includes example Boolean trajectories illustrating one possible behavior for each template. In the following, we use α and β to denote non-temporal subformulas, each being an atomic proposition or a Boolean combination of atomic propositions defined by the grammar in Section 2.3. More complex formulas can be then generated through Boolean combinations and temporal nesting of the template forms below.

**Figure 3.**
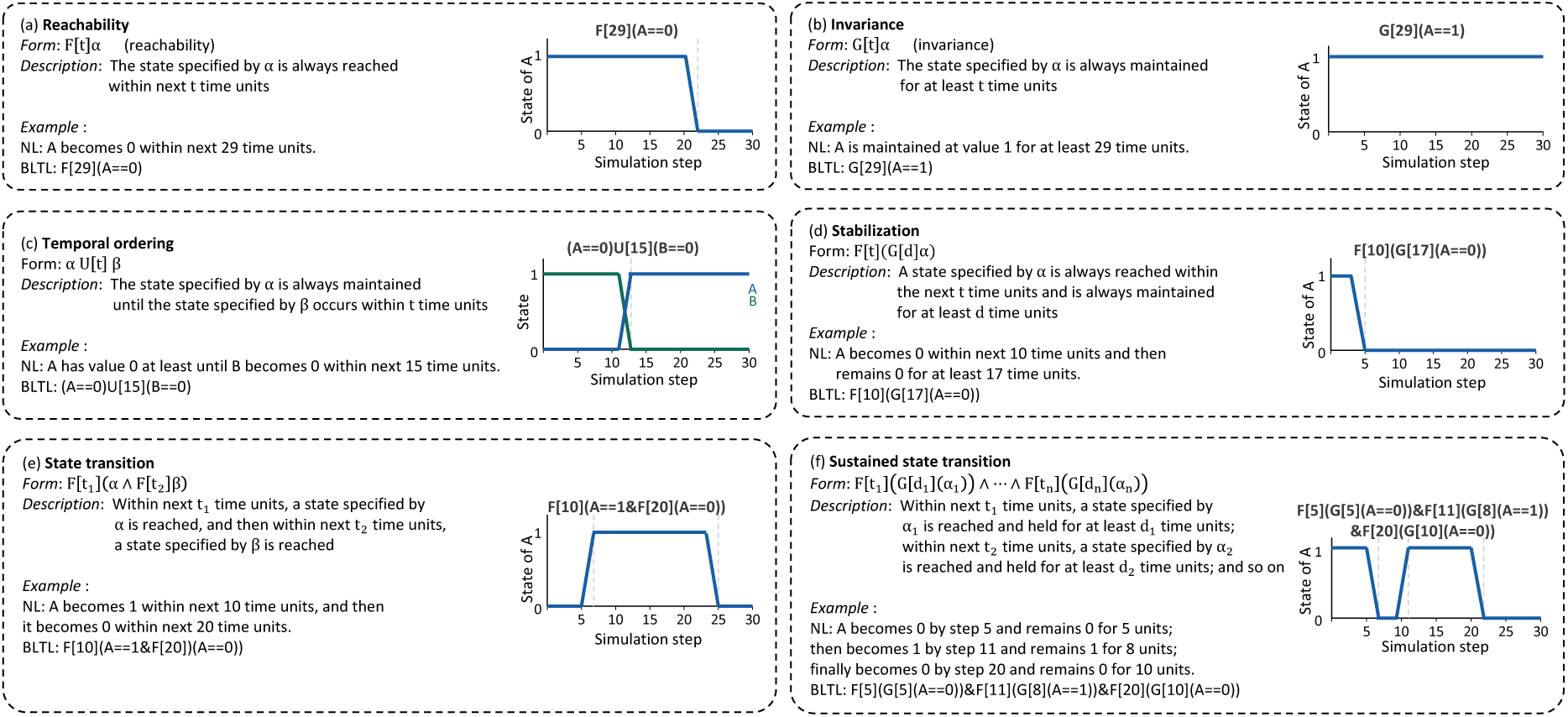
Representative BLTL template categories derived from manually curated specifications. (a). Reachability templates describe states that become true within a bounded time window. **(b).** Invariance templates describe states that are maintained over a bounded time window. **(c).** Temporal ordering templates specify that one state is maintained until another state occurs. **(d).** Stabilization templates describe states that become true and are then maintained for a specified duration. **(e).** State transition templates capture successive state changes without explicit holding-duration requirements. **(f).** Sustained state transition templates capture successive state changes with explicit holding intervals. Symbols *α*, *β* denote non-temporal state subformulas. Each box includes the template form, a brief description, an example NL-BLTL pair, and an illustrative trajectory for one satisfying behavior. Dashed vertical lines mark state-change boundaries in the example trajectories. More complex nested formulas can be generated through Boolean combinations and temporal nesting of these template forms.

**Reachability template** (Figure 3(a)). This template represents the BLTL structure with the form F[t]α, indicating that the state specified by α becomes true within t time units.

**Invariance template** (Figure 3(b)). This template represents the BLTL structure with the form G[t]α, indicating that the state specified by α is maintained continuously for t time units.

**Temporal ordering template** (Figure 3(c)). This template represents the BLTL structure with the form α U[t] β. It captures a temporal ordering relation in which one element’s behavior must be maintained until another event takes place.

**Stabilization template** (Figure 3(d)). This template allows for nested temporal operators, with the typical form F[t](G[d]α) that represents a state becoming true sometime within the next t time units and holding for at least d time units. It is often used to describe a transient change in the value of a biological element, followed by a period of stabilization in the system. Prior work^10^ also includes the pattern G[t](F[d]α) as a “recurrence” template. However, such patterns are not sufficiently represented in our dataset and are therefore omitted in this study.

**State transition template** (Figure 3(e)). This template captures successive state changes without requiring each state to be maintained for an explicit interval. A representative form is F[t_1_](α ∧ F[t_2_]β), where α becomes true within t_1_ time units, and β becomes true within t_2_ time units from the evaluation point where α holds.

**Sustained State transition template** (Figure 3(f)). This template captures successive state changes with explicit hold intervals, and can be expressed as a conjunction F[t_1_]DG[d_1_](α_1_)E ∧ ⋯ ∧ F[t_n_]DG[d_n_](α_n_)E, where each state specified by α_i_ becomes true within its corresponding time bound and is maintained for d_i_time units. To ensure that the intervals are temporally ordered and non-overlapping, the time bounds satisfy t_i+1_ > t_i_ + d_i_, which allows the next interval to begin only after the previous interval has completed its stabilization window.

### 2.5 Synthetic data generation

A key challenge in training language models that translate NL to BLTL in the biological domain is the lack of annotated NL-BLTL pairs. To overcome this limitation, we constructed a large synthetic dataset consisting of BLTL formulas and their corresponding NL descriptions. As illustrated in Figure 4, the construction pipeline consists of three steps: BLTL formula generation, verification with template distribution control, and NL generation via few-shot prompting. We describe each step in the following. The resulting verified dataset is then used to fine-tune the LLMs as described in Section 2.6.

**Figure 4.**
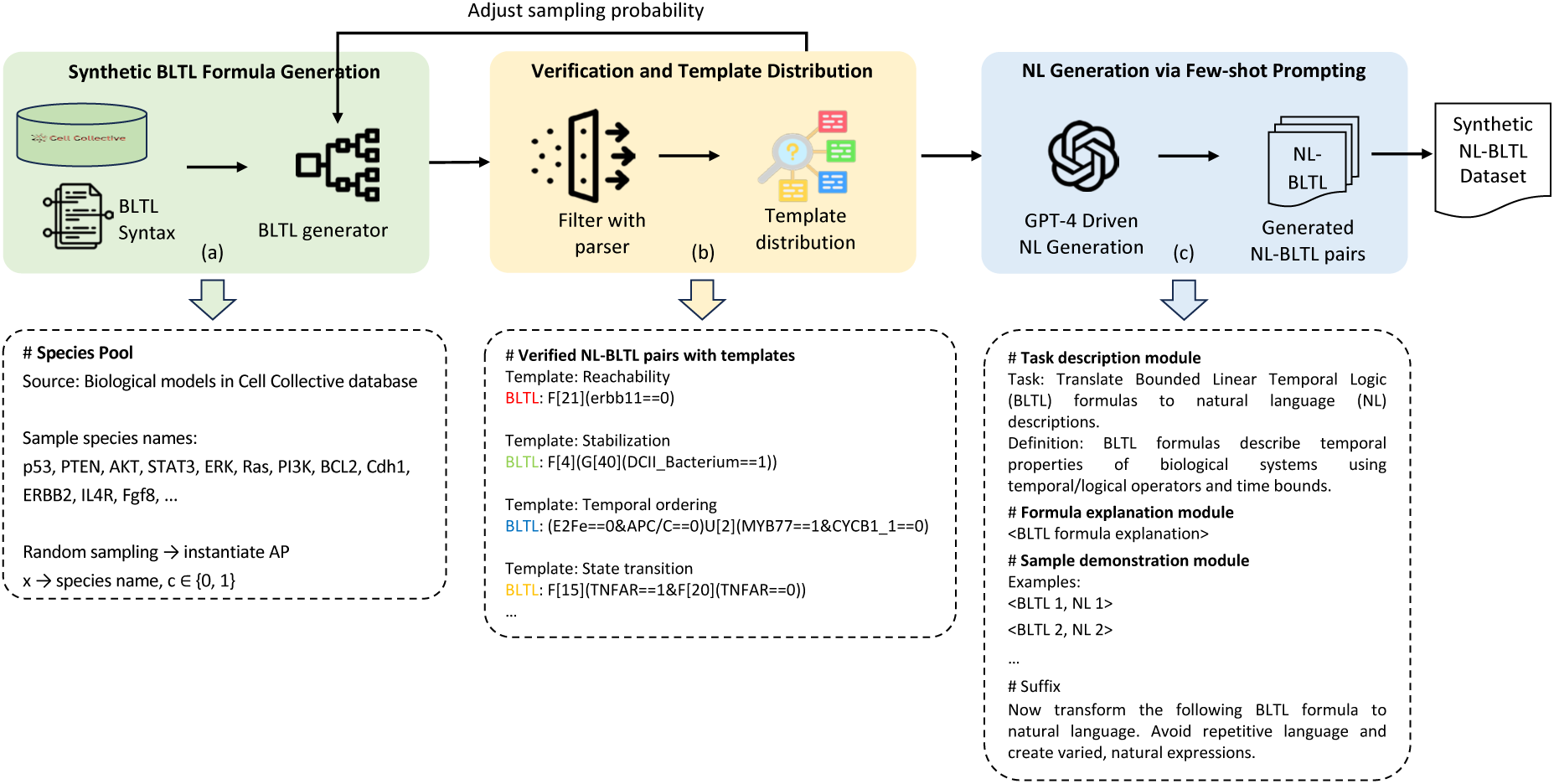
Pipeline of synthetic NL-BLTL dataset construction. (a). The input species names are drawn from 57 curated Boolean models in the Cell Collective database, forming a biologically grounded pool from which the BLTL formula generator samples to instantiate atomic propositions. The grammar that governs the formula structure is shown in Figure 2(a). **(b).** Each synthetic BLTL formula is verified with a parser and assigned to a template category (Section 2.4) to ensure syntactic validity and structural diversity before NL generation. The observed template distribution is used to adjust sampling probabilities for synthetic BLTL formula generation. **(c).** Verified BLTL formulas are passed to GPT-4 using the few-shot prompt template shown here. The prompt includes a Task description module, a Formula explanation module, a Sample demonstration module, and a closing instruction requesting diverse and semantically faithful natural language (NL) descriptions. The resulting verified BLTL formulas and generated NL descriptions form the final synthetic NL-BLTL dataset.

#### 2.5.1 BLTL formula generation

To generate synthetic BLTL formulas (Figure 4(a)), we designed a recursive, tree-based formula generation algorithm summarized in Algorithm 1. Our BLTL generation process combines probabilistic operator sampling with biologically grounded atomic propositions. The algorithm adaptively controls structural complexity by limiting operator depth and reusing variables to maintain temporal consistency across sequential phases. Species names in Cell Collective models are collected, filtered, and normalized to form the species pool S.

##### Algorithm 1: Synthetic BLTL formula Generation

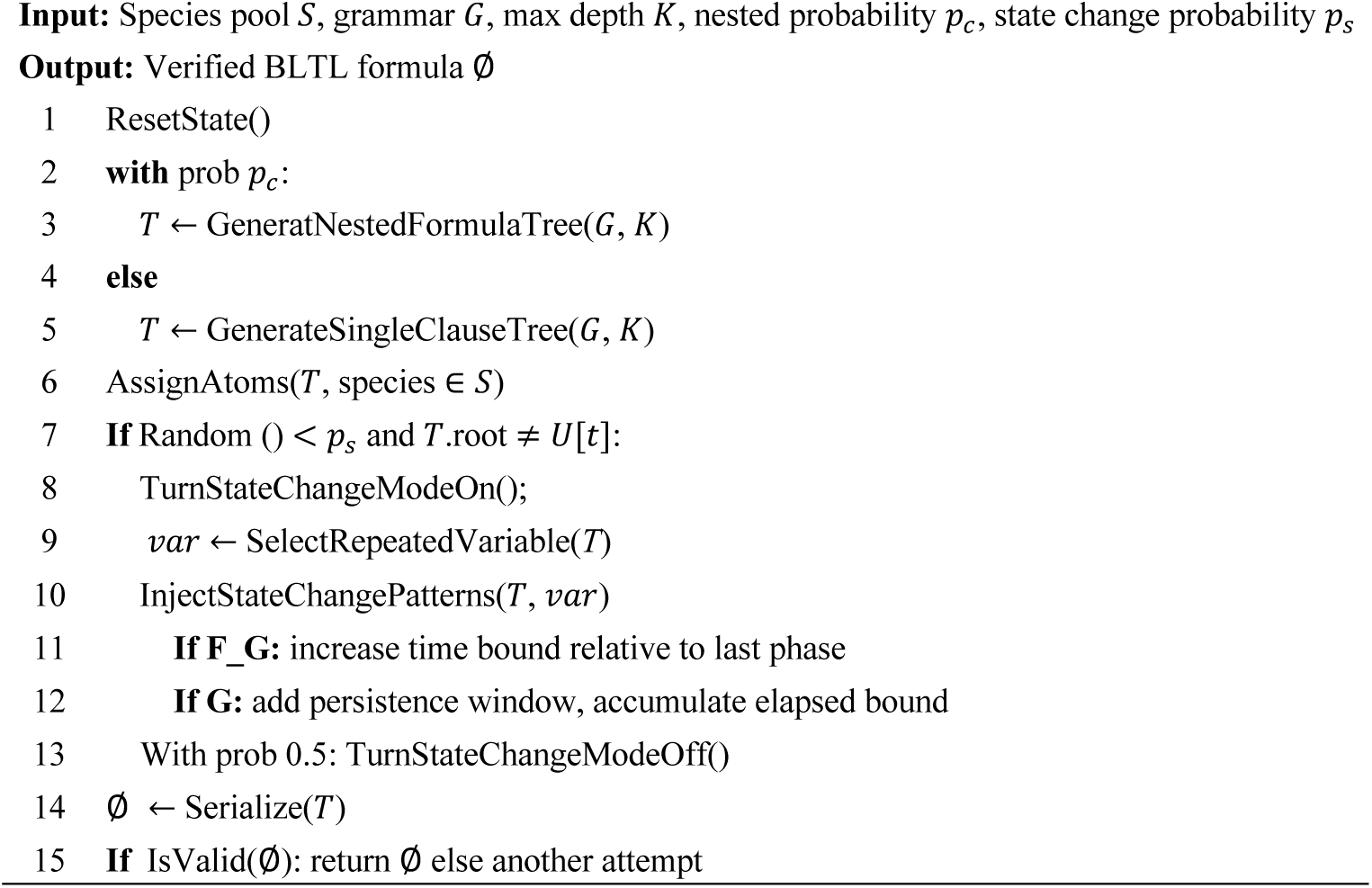

The algorithm begins by sampling a structural template from the grammar defined in Section 2.3. With probability *p_c_*, it generates a nested multi-clause formula (i.e., a Formula node with several Clause children) or a single-clause formula. The grammar is then recursively expanded up to a fixed depth K, producing a structural formula tree T, with internal nodes corresponding to temporal operators F[t], G[t], or the bounded Until operator U[t], and leaves being atomic propositions. After the structural formula tree is created, species variables are instantiated by randomly sampling from the species pool S and assigning each sampled variable a Boolean value. With probability *p_s_*, the algorithm then augments the tree with state-change patterns for a selected variable. Note that when state-change patterns are turned on, the algorithm maintains a global time bound for the selected variable and incrementally increases this bound when constructing outer temporal operators to ensure temporal consistency across phases. The completed formula tree is serialized into a string and passed to the syntax-verification step described in Section 2.5.2, where formulas that fail grammar or parser checks are removed from the final synthetic BLTL set.

#### 2.5.2 Verification and template distribution

Before NL generation, we incorporate BLTL syntax verification and template distribution control to ensure structural validity and diversity across generated BLTL formulas. Each generated BLTL formula is filtered through a strict parser, which checks syntax correctness, validates operator nesting, and confirms that the formula can be converted into a programmable directed acyclic graph (DAG) representation compatible with model checking tools. Only formulas that pass all checks are retained for subsequent NL generation.

To promote structural diversity, we control the template distribution by assigning retained BLTL formulas to template categories and adjusting the sampling probabilities *p_c_* and *p_s_*, as shown in Figure 4(b).

#### 2.5.3 NL generation via few-shot prompting

For each verified BLTL, we employ GPT-4 to generate a semantically aligned NL description through in-context learning. We provide few-shot demonstrations within the prompt to help the LLM capture the correspondence between temporal logic expressions in BLTL and their NL counterparts. The prompt design is shown in Figure 4(b) and consists of three functional modules.

##### Task description module

This module specifies the task as generating NL descriptions from BLTL formulas and frames how a BLTL formula expresses temporal properties of biological systems. A closing instruction asks the model to produce outputs that preserve formula semantics while differing from the demonstrations, ensuring diversity across the generated dataset.

##### Formula explanation module

We construct a brief explanation of the BLTL formula syntax to help the LLM understand the formal structure of the input formulas. It introduces both temporal and logical operators along with their respective symbols and semantic interpretations.

##### Sample demonstration module

This module includes several selected manually labeled BLTL formulas with representative patterns and their corresponding NL descriptions. These demonstrations act as semantic anchors for LLMs’ references so that LLMs can better understand the structure and content of the BLTL formula and how formal temporal logic structures correspond to natural language.

### 2.6 Model fine-tuning

We mainly use Qwen-7B^27^ as the base LLM for fine-tuning on our synthetic NL-BLTL dataset. To study the impact of model size, we also fine-tune three LLMs on the same data: DeepSeek-7B^28^ and Llama3.1- 8B^29^, which have similar model sizes, and BioMistral-7B^30^, a LLM pretrained on large-scale biomedical corpora. Furthermore, we fine-tuned T5-large^14^, a sequence-to-sequence language model, to assess the effectiveness of various model architectures on this generation task.

### 2.7 NL preprocessing with Chain-of-Thought method

Although the generated synthetic dataset (Section 2.5) provides semantically valid NL-BLTL pairs for training the LLMs, real-world NL descriptions derived from hypotheses about biological system behaviors are often ambiguous in their temporal expressions. To enhance the interpretability and temporal accuracy of NL inputs and further improve the accuracy of BLTL generation, we apply the CoT method to the input NL descriptions prior to BLTL generation. Specifically, we design a NL preprocessing approach based on the CoT method, which decomposes each input sentence into intermediate reasoning steps that clarify its temporal and logical structure and perform necessary rephrasing before translating them into BLTL formulas. Through this process, implicit or composite temporal relations in biological text can be clarified to ensure the inputs are aligned with the formal semantics of BLTL.

We first design a set of instructions that the LLMs should follow during the generation. For example, the model is instructed to extract time bounds by identifying numerical time references (e.g., 10 from "by round 10", 20 from "within 20 time units"), and to detect whether there is a sequence of value changes over time in each variable. To adapt the CoT method to diverse biological phrasing styles and temporal scales, we append few-shot examples in the prompt that illustrate the desired step-by-step reasoning for temporal and logical decomposition, guiding LLM’s understanding and preprocessing of complex natural language inputs. The preprocessing step therefore produces both a rephrased NL description and a set of extracted constraints (e.g., biological elements and time bounds). The rephrased NL descriptions are then passed to the fine-tuned LLMs for BLTL generation described in Section 2.8.

### 2.8 BLTL generation with grammar-constrained decoding

Following the CoT-based preprocessing step described above, we employ the GCD strategy for constraining the outputs of the fine-tuned LLMs to the defined BLTL_R_ formulas (Formula). The preprocessing step resolves linguistic ambiguity and extracts key information (e.g., biological elements, explicit time bounds), which we incorporate into the context-free grammar used for decoding. As stated earlier in Section 2.3, the predefined BLTL_R_ specifications are encoded into this grammar, where the extracted constraints are injected to restrict allowable biological elements and time bounds to those identified during preprocessing. During decoding, each token generated by the fine-tuned LLMs must satisfy the grammar rules. This restricts the search space and guides the model toward generating syntactically valid BLTL_R_ formulas. Given the recursive and nested structure of BLTL_R_ specifications, GCD is particularly suitable for enforcing well-formed outputs in our settings. In addition, the integration of GCD also mitigates hallucination of LLMs, which may otherwise generate biological elements that are not present in the input sentence or produce incorrect time bounds.

## 3 Experimental setup

### 3.1 Datasets

We conducted experiments on two NL-BLTL datasets: a synthetic dataset and a manually curated biomedical dataset. For the biomedical dataset, we curated 97 NL-BLTL pairs by extracting temporal properties and crafting corresponding NL descriptions based on a previous study on naïve T cells^26,31^. We additionally collected 20 NL-BLTL pairs from a related work on PCC microenvironment^25^. These two sets are concatenated to form the biomedical NL-BLTL dataset, which serves as the gold-standard dataset for evaluating BLTL generation in biological contexts. Following the data generation pipeline described in Section 2.5, we constructed a synthetic dataset of 5,000 NL-BLTL pairs, with species names drawn from 57 biological models manually downloaded from the Cell Collective database. The dataset is split into 3,000 samples for training and 1,000 samples each for validation and testing.

While the BLTL_R_ specifications are generally applicable to a broader range of values, for our experiments we used Boolean biological models, in which each species takes a binary activation state. The binary abstraction forms the basis of most prior BLTL verification studies^25,26^, has been broadly used in systems biology ^31–34^, and matches the representation of the curated models from the Cell Collective database that we used in this study. Accordingly, in atomic propositions we restrict to c ∈ {0, 1}, representing the Boolean activation state of a species.

### 3.2 Evaluation metrics

In our NL-BLTL task, the objective is to generate structured BLTL formulas from NL that depict the temporal behaviors of specific system elements in the domain of systems biology. Essentially, this task can be framed as a text generation problem, where the output is a BLTL formula generated by LLMs. To evaluate the performance of the models in generating BLTL, we adopted multiple metrics that assess both syntactical and semantic correctness. The evaluation metrics are described below.

**BLEU score.** The BLEU (Bilingual Evaluation Understudy)^35^ is a common metric used to evaluate the quality of machine-translated text by comparing it to the reference translations. Specifically, BLEU compares consecutive sequences of words (n-grams) between generated and reference outputs.

**Exact Match (EM).** A straightforward metric that assesses how many generated outputs exactly match the reference outputs. Additionally, we adopt a strategy to make EM more tolerant. Specifically, if the predicted and gold BLTL formulas are not exactly matched, they will be parsed into DAG-style logic representation by the syntax verification module, and an EM match is counted if their representations are equivalent.

**Validity.** This metric evaluates the structural validity of BLTL formulas. We calculate it as the proportion of generated outputs that pass the verification step described in Section 2.5.2.

### 3.3 Implementation

We conducted the comparison experiments using three categories of models for BLTL generation, including an out-of-the-box LLM (GPT-4^21^), open-source decoder-only LLMs (Qwen2.5-7B-Instruct^36^, deepseek-llm-7b-chat^37^, Llama-3.1-8B^38^, and BioMistral-7B^39^) and an encoder-decoder baseline (T5- large^40^). As LLMs are pretrained on large-scale text corpora containing scientific and logical content, we aim to examine their inherent generalization capability for NL-BLTL generation. To assess this capability, all models are trained exclusively on the synthetic NL-BLTL dataset, using a 6:2:2 split for training, validation, and testing. The biomedical NL-BLTL dataset is used solely for evaluation, assessing whether BLTL patterns learned from synthetic data transfer to real-world biological applications.

Due to limited computing resources, we apply instruction tuning using LoRA^41^ for parameter-efficient fine-tuning of the open-source LLMs, which updates only the low-rank adapter layers while keeping the base model weights frozen. T5-large model is trained through full-parameter supervised learning. For the CoT-based NL preprocessing and GCD experiments, we only apply these techniques to models for which they are compatible. Among the open-source LLMs, we select Qwen2.5-7B as the backbone as it outperformed other models in our preliminary evaluations. GPT-4 is also tested with CoT prompting. However, GCD cannot be applied to GPT models, as their APIs do not expose token-level logits or allow custom decoding constraints. Likewise, T5-large does not benefit from CoT and GCD, since both techniques rely on the autoregressive decoding, which is incompatible with T5’s encoder-decoder architecture. All finetuning runs use a learning rate of 1e-4, batch size of 8, and 10 training epochs. Experiments are conducted on a server using an Nvidia A100 GPU. GPT-4 baselines are evaluated using zero-shot, CoT-augmented, and few-shot prompting without any parameter updates. In the zero-shot setting, GPT-4 is instructed to generate BLTL formulas using a structured system prompt. For the few-shot setting, the demonstrations are sampled from our synthetic NL-BLTL training set.

## 4 Results

### 4.1 Performance on the synthetic NL-BLTL dataset

Table 1 reports model performance on the synthetic NL-BLTL test set, which evaluates a model’s ability to recover compositional BLTL structure under idealized alignment conditions. Note that CoT preprocessing and GCD are not applied in this experiment, as the synthetic NL-BLTL pairs are automatically generated and already aligned in structure. GPT-4 with zero-shot and few-shot prompting achieved moderate EM scores (42%-44%) and BLEU score around 82%, indicating that powerful out-of-the-box LLMs can partially reconstruct BLTL structure. The marginal improvement for GPT-4 by adding few-shot demonstrations suggests that prompting alone cannot reliably teach the LLMs complex BLTL syntax, such as nested temporal operators, without explicit supervision. T5-large model and open-source LLMs achieve substantially higher performance after fine-tuning with the synthetic data. T5-large obtains an EM score of 82.90%, demonstrating the effectiveness of directly supervised learning for the NL-BLTL mapping. Despite being pretrained on biomedical corpora, BioMistral does not surpass the other open-source LLMs after fine-tuning, suggesting that direct BLTL generation from an aligned NL does not require much domain knowledge. Qwen2.5 achieves the highest BLEU (97.08%), EM (83.50%) and Validity (100.00%) scores among all evaluated models, outperforming other open-source LLMs including DeepSeek, Llama3, and BioMistral. This demonstrates Qwen’s superior robustness in BLTL generation. Overall, these results confirm that fine-tuning on synthetic data is both necessary and sufficient for strong performance under idealized conditions, and that the choice of foundation model meaningfully affects outcomes.

**Table 1:** Evaluation results on the synthetic NL-BLTL dataset.

| Model | Synthetic NL-BLTL |  |  |
| --- | --- | --- | --- |
|  | BLEU | EM | Validity |
| GPT-4o (Zero-shot) | 81.53 | 42.70 | 98.70 |
| GPT-4o + Few-shot | 82.51 | 44.00 | 95.60 |
| T5-large | 97.07 | 82.90 | 99.90 |
| BioMistral | 96.83 | 82.30 | 98.80 |
| DeepSeek | 96.42 | 81.30 | 99.50 |
| Llama3 | 90.25 | 77.3 | 96.80 |
| Qwen2.5 | **97.08** | **83.50** | **100.00** |

### 4.2 Performance on the biomedical NL-BLTL dataset

Table 2 reports performance on the biomedical NL-BLTL dataset, which presents a more challenging evaluation setting due to human-annotated descriptions with diverse phrasings, unseen entities, and implicit temporal semantics. These experiments assess whether models trained exclusively on the synthetic dataset can generate correct BLTL formulas from NL descriptions in a different biological context. As described earlier, we decompose the NL-BLTL task into two subtasks: CoT prompting with GPT-4 to rewrite ambiguous NL expressions and extract biological variables and time bounds, followed by BLTL generation with finetuned LLMs using GCD, where the extracted variables and time bounds are embedded in the decoding grammar.

**Table 2:** Evaluation results of multiple methods on the biomedical NL-BLTL dataset.

| Model | BLEU | EM | Validity |
| --- | --- | --- | --- |
| GPT-4o (Zero-shot) | 78.43 | 41.03 | 98.29 |
| GPT-4o + CoT | 83.36 | 49.57 | 99.15 |
| GPT-4o + Few-shot | 61.41 | 64.96 | 89.74 |
| GPT-4o + Few-shot + CoT | 70.69 | 68.38 | 94.02 |
| T5-large | 65.36 | 22.22 | 52.14 |
| Qwen2.5 + CoT + GCD (ours) | **91.34** | **84.62** | **100.00** |

The results show that combining CoT preprocessing and GCD performs the best. Across all three metrics, our method achieves the highest scores of 91.34% for BLEU, 84.62% for EM and 100.00% for Validity, demonstrating strong generalizability to real biomedical language descriptions. GPT-4 with few-shot prompting and CoT achieves 68.38% EM, more than 16 points below our method, confirming that fine-tuning on synthetic data provides critical structural knowledge that prompting alone cannot replicate. T5- large performs poorly on this dataset (22.22% EM, 52.14% Validity), indicating that encoder-decoder architectures struggle to generalize beyond their training distribution to the linguistic diversity of real biomedical text.

### 4.3 Ablation study

Table 3 presents an ablation study on the biomedical dataset quantifying the individual and combined contributions of CoT preprocessing and GCD. The base fine-tuned Qwen2.5 model without either component achieves 70.09% EM and 99.15% Validity. Adding CoT alone improves EM to 82.05%, confirming that resolving temporal ambiguity in biological NL before translation is a critical factor in biomedical generalization. Adding GCD alone improves EM to 77.78% and raises Validity to 100.00%, demonstrating that grammar constraints effectively prevent hallucinated variables and invalid formula structures. The combination of CoT and GCD achieves the best performance at 84.62% EM and 100.00% Validity, a 14-point gain over the base model, confirming that the two components address complementary failure modes, semantic ambiguity and syntactic invalidity, respectively, and that their benefits are additive rather than redundant.

**Table 3:** Ablation study on the biomedical NL-BLTL dataset. We evaluate the contribution of NL preprocessing with CoT and GCD for BLTL generation.

| Model | BLEU | EM | Validity |
| --- | --- | --- | --- |
| Qwen2.5 | 84.54 | 70.09 | 99.15 |
| Qwen2.5 + CoT | 87.65 | 82.05 | 98.29 |
| Qwen2.5 + GCD | 87.53 | 77.78 | 100.00 |
| Qwen2.5 + CoT + GCD (ours) | **91.34** | **84.62** | **100.00** |

### 4.4 Scaling effect

Figure 5 presents scaling effect results across model variants on both datasets as the number of synthetic training examples increases from 100 to 3,000. On the synthetic test set, performance improves consistently with training data size, reaching a maximum EM of 83.50% at 3,000 examples. On the biomedical dataset, all models except T5-large show substantial improvement as training data grows from 100 to 500 examples, after which gains become more gradual. Fine-tuned Qwen2.5 rises from 53.85% to 70.09% EM across this range before reaching 84.62% with CoT and GCD. These results indicate that pretrained LLMs are data-efficient learners of NL-BLTL mappings, benefiting quickly from relatively small amounts of synthetic training data, while continued improvements at larger scales confirm that task-specific fine-tuning remains essential for optimal performance.

**Figure 5.**
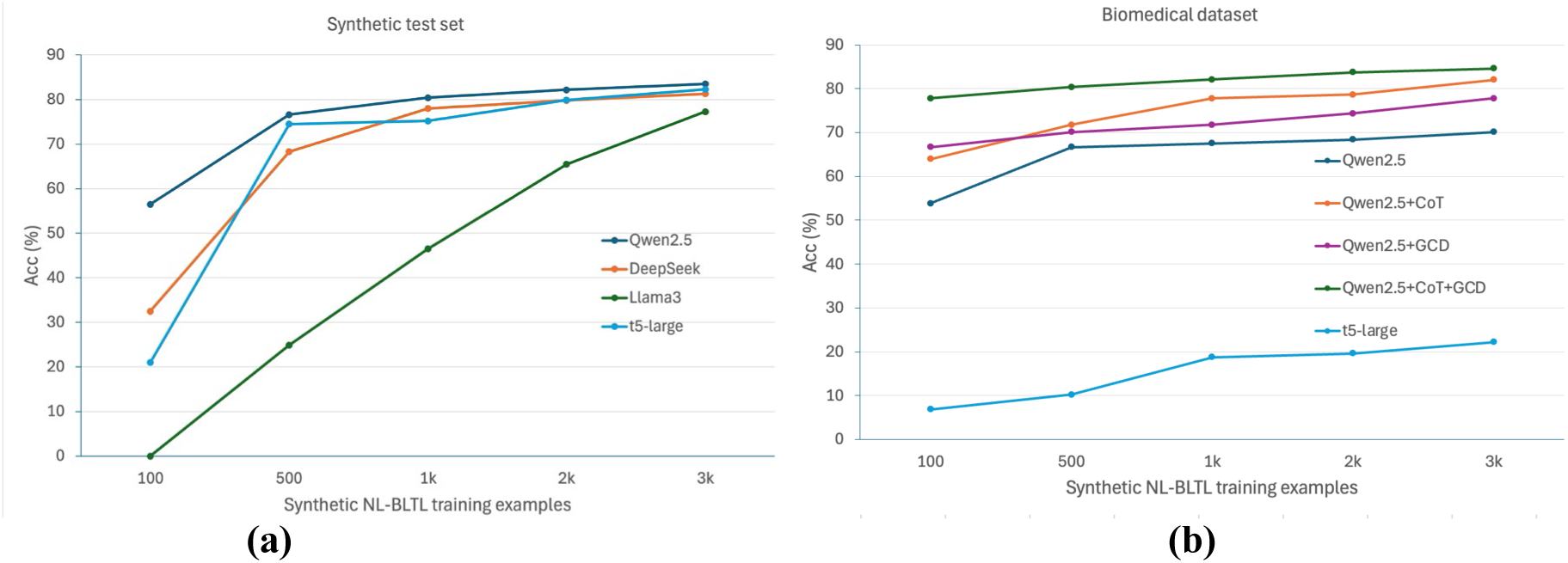
**Scaling effect of synthetic training data on model accuracy for (a) the synthetic test set and (b) the biomedical dataset.**

### 4.5 Error analysis

Table 4 presents three representative error categories identified from incorrect predictions of our best model on the biomedical dataset. Temporal operator mismatches occur when subtle persistence or precedence cues such as "maintain … until" are mapped to incorrect operators, as in Example 1 where the until relation (p52==0)U[12000](apoptosis==1) is predicted as F[12000](G[12000](apoptosis==0&p53==0)), losing its precedence semantics. Temporal scope mismatches arise when independently bounded behaviors of multiple elements are collapsed into a single operator scope, producing a strictly stronger formula, as illustrated in Example 2. Finally, proposition-level errors emerge when the model inserts atomic propositions or subformulas that are not supported by the input or are semantically inconsistent with it. In Example 3, the model adds pten==0 although the input indicates that pten should be 1. These error categories highlight three directions for future improvement: richer training coverage of persistence and until patterns, explicit supervision of operator scope for multi-element descriptions, and consistency constraints on subformulas within a BLTL specification.

**Table 4:**
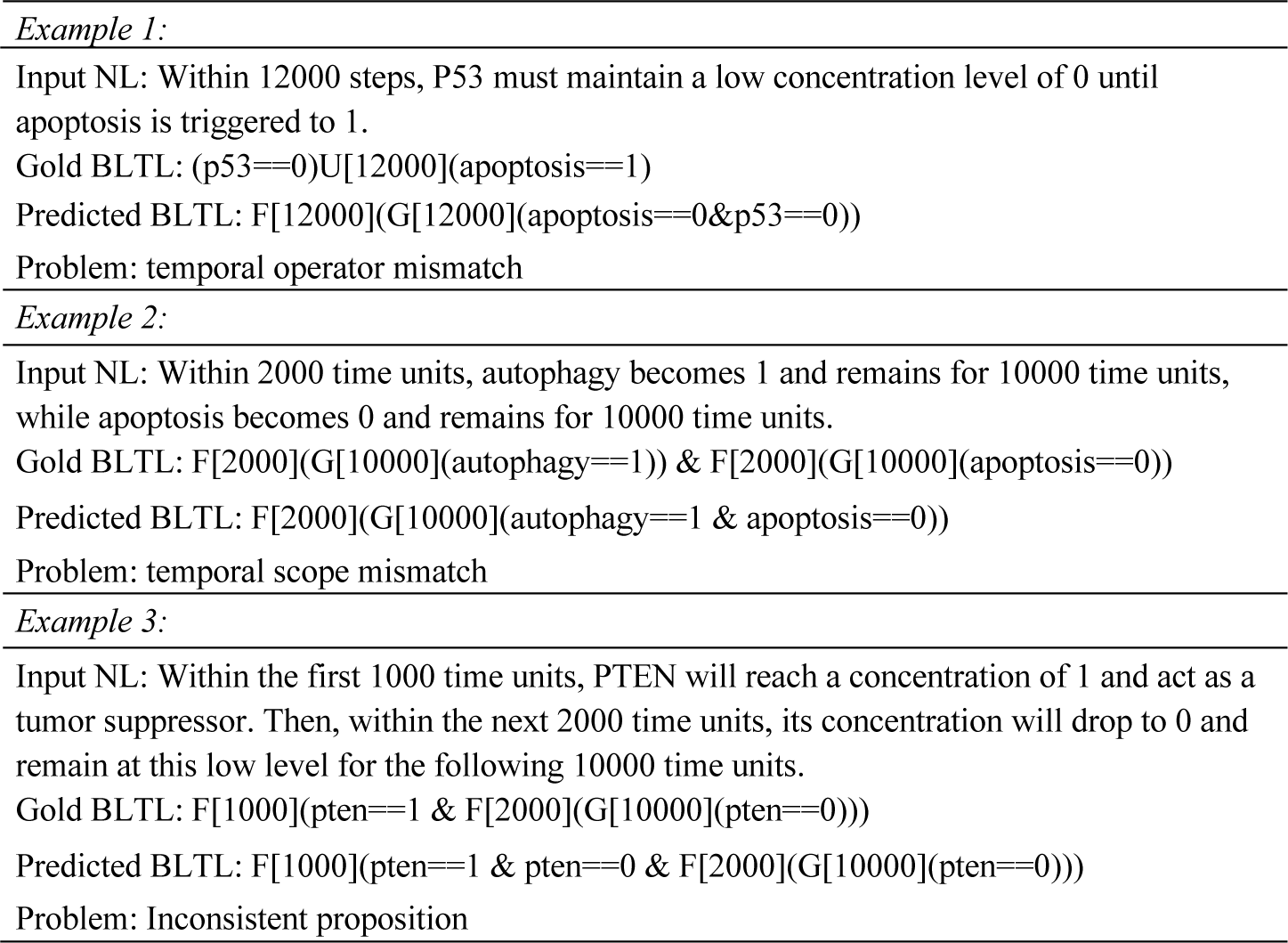
Error examples for evaluating the best model on the biomedical NL-BLTL dataset.

## 5 Discussion

NL2BLTL demonstrates that automated translation of natural language biological propositions into valid BLTL formulas is not only feasible but practically achievable at high accuracy, with direct implications for the integration of formal verification into routine systems biology workflows. By making statistical model checking accessible without requiring biologists to learn temporal logic syntax, NL2BLTL removes a longstanding bottleneck in the model verification pipeline and opens new possibilities for systematic, large-scale hypothesis evaluation against computational models of biological systems.

A key finding across both datasets is that fine-tuned open-source LLMs substantially outperform GPT-4 under all prompting configurations, confirming that the structural complexity of BLTL, particularly nested temporal operators and strict variable alignment, cannot be reliably learned through prompting alone and requires explicit supervision on domain-specific data. This observation is consistent with broader biomedical natural language processing evaluations showing that task-specific fine-tuning remains important for reliable LLM performance^42^. The superior performance of Qwen2.5 over other comparably sized models further highlights that foundation model selection is a meaningful design decision for structured biological language generation tasks, and that general biomedical pretraining, as in BioMistral, does not confer an advantage when the target output is a formal logical structure rather than natural language. In practice, models achieving high BLEU but lower EM may be suitable for semi-automated BLTL drafting with human curation, while models achieving high EM and perfect Validity, as demonstrated by our full framework, are reliable for fully automated downstream use in simulation and model checking.

Several limitations of the current work point toward important directions for future development. First, the synthetic NL descriptions, while structurally diverse, do not fully capture the linguistic variety of real biomedical narratives, including passive constructions, hedged language, and cross-sentence temporal dependencies, suggesting that expanding the manually curated biomedical dataset and incorporating it into fine-tuning would improve robustness in real-world deployment. Second, the current CoT preprocessing step relies on GPT-4 with manually designed prompts and few-shot examples, which may not scale efficiently to new biological domains or temporal logic templates without prompt revision. Automated information extraction pipelines, including biomedical entity normalization^43^ and retrieval-augmented biomedical LLMs^44^, could help identify and align biological entities, temporal cues, and relational operators directly from text would substantially enhance the scalability and generalizability of the framework. Extending NL2BLTL beyond Boolean models to handle multi-valued or continuous biological variables, and complementing exact match evaluation with semantic equivalence checking, since logically equivalent formulas can differ structurally, are additional priorities for future work. Despite these limitations, the flexibility of the CoT and GCD components, whose constraints are derived directly from preprocessing outputs, ensures that NL2BLTL can adapt to new temporal logic templates or biological domains without architectural modification, providing a strong foundation for continued development.

## Acknowledgements

This work was funded in part by the NSF EAGER award CCF-2324742, the NSF CAREER award CCF- 2442884, and the NIH NLM award R01LM014673. This research was supported in part by the University of Pittsburgh Center for Research Computing, RRID:SCR_022735, through the resources provided. Specifically, this work used the H2P cluster, which is supported by NSF award number OAC-2117681.

